# Integrating phylogenetic and functional data in microbiome studies

**DOI:** 10.1101/2022.02.21.480893

**Authors:** Gavin M. Douglas, Molly G. Hayes, Morgan G. I. Langille, Elhanan Borenstein

## Abstract

Microbiome functional data are frequently analyzed to identify associations between microbial gene families and sample groups of interest. This is most often performed with approaches focused on the metagenome-wide relative abundance of microbial functions. Although such approaches can provide valuable insights, it is challenging to distinguish between different possible explanations for variation in community-wide functional profiles by considering functions alone. To help address this problem, we have developed a novel, phylogeny-aware framework to expand taxonomic balance tree-based approaches to identify enriched functions more robustly. The key focus of our approach, termed POMS, is on identifying functions that are consistently enriched in sample groups across independent taxonomic lineages. Based on simulated data we demonstrate that POMS can more accurately identify gene families that confer a selective advantage compared with commonly used differential abundance approaches. We also show that POMS can identify enriched functions in real-world metagenomics datasets that are potential targets of strong selection on multiple members of the microbiome. While this framework may not be able to identify all potential functional enrichments, the enrichments it does identify are more interpretable and conservative compared with those identified by existing differential abundance approaches. More generally, POMS is a novel approach for exploring microbiome functional data, which could be used to complement standard analyses. POMS is freely available as an R package at: https://github.com/gavinmdouglas/POMS.

## Introduction

Microbiome sequencing has been applied to characterize myriad environments and is typically analyzed based on the relative abundance of microbial features. These features may include both taxa (the microbes present) and functions (the genes and pathways they encode). While both data types have been leveraged to make valuable observations, they are typically analyzed independently. Yet, linking these data types is clearly required to make coherent interpretations of observed shifts in microbiome data^1,2^. For example, an enrichment of a particular microbial pathway in a group of samples (e.g., a disease or unique environment) could represent selection on multiple independent taxa which possess that pathway, which would be of significant biological interest. In contrast, such an enrichment could have also arisen due to the presence of that pathway in a specific microbial taxon that is more abundant for various other reasons in the group of interest, making this enrichment less biologically interesting.

Two approaches have been developed that partially address the challenge of how to link taxonomic and functional data types. FishTaco^2^ is a tool that pinpoints the taxonomic contributors to functional shifts identified by differential abundance tests. FishTaco, however, is a post-hoc tool that is applied *after* significant microbial functions have been identified, whereas ideally, functional and taxonomic data would be integrated *while* testing for differential functions to better identify strong enrichment candidates.

Phylogenize is an approach that explicitly integrates functional and taxonomic data during statistical testing^3,4^. It identifies functional associations based on the prevalence of taxa that encode each gene family. This is performed using phylogenetic linear models, which account for the genetic similarity of co-occurring taxa that arises due to their shared evolutionary history. Using this framework, significant gene families and pathways that are contributed by a diverse set of taxa from within a given phylum are identified. Although this approach forms a key improvement over past analytical approaches, it also has some limitations. Most notably, it is intended to be used with a pre-existing database^5^, which means that novel genomes from metagenomics sequencing datasets cannot be used as input. It also focuses solely on the presence/absence of gene families, meaning that it does not incorporate taxon relative abundances. Last, phylogenize restricts its analyses only to taxa within a single phylum at a time, which means that key functions distributed across multiple phyla could be overlooked. For these reasons, phylogenize cannot be appropriately compared with standard differential abundance approaches applied to microbiome data, and continued development of novel methods for joint taxonomic and functional data analysis is needed.

Some of the complexity of integrating taxonomic and functional analyses, as well as other challenges in microbiome data analysis, stem from the difficulties of applying standard statistical approaches to raw microbiome data due to their compositional nature. Fortunately, there is growing interest in improved compositional approaches for analyzing microbiome data. Specifically, analyzing ratios of microbiome feature relative abundances (rather than the abundances of features themselves) has recently been proposed as a solution to the compositionality problem^6,7^. However, it is unclear which features should be used for stably computing these ratios. One proposed solution is to compare ratios of taxa (based on the isometric log-ratio transformation) on each side of every node in a phylogenetic tree that links the various taxa^8^. This general approach is now commonly referred to as analyzing balance trees^9^. While this is a statistically valid framework for analyzing microbiome data, it is often unclear how to interpret differences in taxonomic ratios.

Herein, we aim to address both the challenges involved in functional differential abundance analysis and the limited interpretability of balance tree-based approaches, by testing for functional enrichment in a balance tree framework. Specifically, our approach, termed Phylogenetic Organization of Metagenomic Signals (POMS), focuses on identifying cases where multiple taxa that encode a given function are consistently associated with a sample group. Such cases provide more support to the hypothesis that the function itself confers a selective advantage to microbes in the relevant sample group, rather than that the function happens to be present in just a few taxa that are more abundant in that group. We show that this approach can pinpoint interesting functional enrichments in both simulated and real metagenomics data, providing a valuable proof-of-concept that integrating functional enrichments into balance tree analyses improves their interpretability and provides novel insights. Importantly, however, since POMS cannot identify functions with limited taxonomic breadth as highly enriched, we see it not as a replacement for current approaches, but as a complementary tool.

## Results

POMS is a balance tree framework (implemented as an R package) for analyzing microbial functions (**Figure 1**), including both gene families and higher-level functions. The key input tables correspond to taxonomic abundances across samples and per-taxon functional abundances. A phylogenetic tree linking all the taxa present in the samples must also be provided. The per-taxon functional abundance table corresponds to genome annotations for metagenome-assembled genomes (MAGs), or other known taxa present in an environment. This format contrasts with the functional abundance tables that are largely unlinked from specific taxa, such as those that are produced through read mapping against a database of broadly distributed gene families. The key output of POMS is a table summarizing, for each annotated function, the results of the test for consistent enrichment. Information regarding the intermediate steps of the workflow (described below) is also provided.

**Figure 1:**
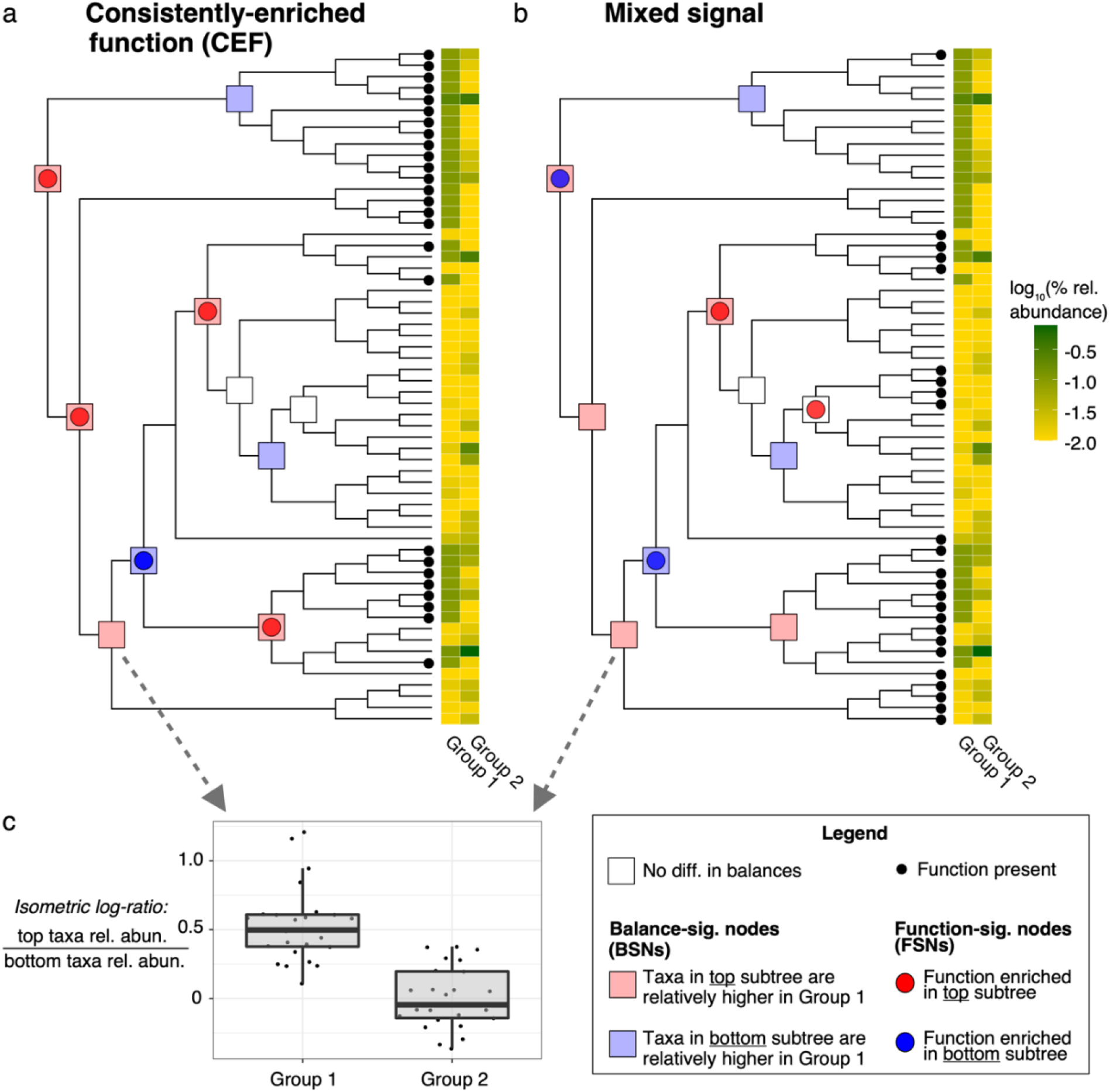
Contrasting examples that illustrate the POMS methodology. (a and b) The two phylogenetic trees correspond to 60 microbial genomes found across two sample groups (Groups 1 and 2). The mean log10 relative abundance of these genomes in each group is indicated by the heatmaps. The squares indicated on tested nodes indicate whether there is a difference in the isometric log-ratio of the subtree taxonomic relative abundances between the two sample groups. Red and blue squares are coloured to indicate which subtree is at relatively higher levels in Group 1 samples (relative to the other subtree; see panel c for an example). Note that the taxonomy-based annotations are identical between the two panels. Panels a and b show the distribution of two different microbial functions across these genomes (small black dots at the tips of trees). The coloured circles on the tested nodes indicate that the function of interest is enriched in one subtree compared with the other. POMS tests whether the intersection between coloured squares and circles is higher than expected by chance. Functions that are consistently enriched in the sample group direction (i.e., where the circle and square colours match at nodes consistently, as in panel a, or alternatively where they consistently mismatch) are particularly of interest. Panel b is a contrasting example where the function is not consistently enriched in taxa relatively higher in the same sample group. Note that the minimum number of underlying tips on each side for a node to be retained for analysis was set to four for this example, but it would normally be higher.

The workflow begins by first identifying all nodes in the tree with sufficient underlying tips (10 by default) on both the left- and right-hand sides. All nodes that do not fit these criteria are excluded from the analysis. The balances of taxa (per sample) at the remaining nodes, defined as the isometric log-ratios of the relative abundances of taxa on the left-hand side compared with those on the right-hand side, are then computed. Standard statistical tests can then be used to determine whether these phylogenetic balances differ between sample groups. Put simply, a node with significantly different balance between sample groups indicates that the taxa on either the left- or right-hand side of that node are at significantly higher abundances relative to the taxa on the other side of the node in one sample group compared with the other. By default, significantly different balances between sample groups are identified based on Wilcoxon rank-sum tests. Note that alternatively the user can specify which nodes are significantly different based on an external, user-defined test, which enables more customized analyses.

We refer to each node with a significantly different balance as a *balance-significant node* (BSN). After identifying each BSN, POMS further determines which side of the node is enriched within each sample group. In other words, POMS determines which taxa are at relatively higher levels (relative to taxa on the other side of the node) in each sample group. Notably, such taxa are not necessarily at higher relative abundances in one group or the other, but rather the relative ratios of the taxa on each side of the node are different between the sample groups.

Next, POMS tests for enrichment of each annotated function across the nodes. More specifically, a Fisher’s exact test is computed based on the counts of tips that either do or do not encode the function on either side of the node. Importantly, these enrichment tests are computed at all nodes tested during the balance tree step: not just at BSNs. We refer to each significant node based on this approach as a *function-significant node* (FSN). Again, put simply, an FSN indicates that the tested function is enriched in taxa on either side of the node compared with the other.

Finally, a multinomial exact test is applied to test whether FSNs coincide with BSNs, and consistently in the direction of the same sample group, more often than expected by chance. Specifically, for each tested function, the set of identified FSNs for that function (i.e., nodes at which this function is enriched in one side of the node compared to the other) can be partitioned into three classes. The first class corresponds to FSNs that do not intersect with BSNs (i.e., there is a significant enrichment of a function on one side of the node, but there is no significant difference in sample balances). The other two classes correspond to FSNs that intersect with BSNs: one class where the function is enriched in taxa that are relatively more abundant in the first sample group, and the other class where the function is enriched in taxa that are relatively more abundant in the second sample group. In our example in Figure 1a these two classes are represented by intersecting circles and squares of the same and different colours, respectively. For each function, the multinomial exact test is performed based on the number of FSNs in each class compared with the expected proportions given random intersections between FSNs and BSNs. We refer to significant functions based on this test as *consistently enriched functions* (CEFs).

Although this test is primarily intended to identify CEFs enriched in the direction of a single sample group, this framework can also identify CEFs that show mixed signal of enrichment towards both sample groups. In other words, a function could be significant because there is a depletion of FSNs that do not intersect with BSNs compared with the random expectation, and not that it is consistently enriched towards a particular sample group. Such cases could still be biologically interesting, but this highlights that post-hoc consideration of the number of FSNs of each class is needed to be able to interpret CEFs appropriately.

To validate POMS, we first generated simulated datasets based on samples containing MAGs. The MAGs from shotgun metagenomics (MGS) control samples, with corresponding relative abundances and phylogenetic tree, were obtained from a large human gut MGS meta-analysis^5^. These MAGs had previously been annotated with KEGG orthologs (KOs)^10^. We subsampled 704 control samples into two equally sized groups 1,000 times to create random test datasets.

We then conducted several sets of simulations to evaluate the performance of POMS. In some of these tests we also evaluated the performance of several standard differential abundance approaches: ALDEx2^11^, DESeq2^12^, limma-voom^13,14^, and Wilcoxon tests (based on raw relative abundances or corrected by the abundance of universal single-copy genes [USCGs]). We applied these standard differential abundance approaches to KO abundance tables that lacked taxonomic links. The per-sample abundance of each KO in these tables was computed by summing the product of the taxon abundance and KO copy number for each taxon that encoded the KO.

To characterize the baseline tool behaviour, we first investigated the proportion of significant KOs (based on Benjamini-Hochberg corrected p-values [BH] < 0.05) identified by each tool across the 1,000 random datasets. Most tools identified significant KOs only in a small number of datasets; both POMS and ALDEx2 identified significant KOs in seven datasets, while the relative abundance and USCG-corrected Wilcoxon test approaches identified 17 and 19 datasets with significant KOs, respectively. In contrast, DESeq2 and limma-voom identified significant KOs in 543 and 1,000 datasets, respectively.

We then introduced taxonomic variation between the two groups in each dataset using two different approaches. In the first approach, we randomly selected for each of the 1,000 random datasets, a KO encoded by at least five MAGs. This gene is referred to as the focal gene per dataset and represents a gene that confers a selective advantage in one of the two sample groups to taxa that encode it. To simulate selection acting upon the genomes encoding this gene, we added a pseudocount of one to all genomes that encode the gene and then multiplied the relative abundance of these genomes by 1.5 in one sample group only (**Supp. Figure 1**). This set of simulated datasets will be referred to below as the ‘focal gene’ profiles. In the second approach we conducted analogous simulations, but where random genomes (rather than genomes that encode a specific focal gene) were selected to increase in abundance in one group. For consistency, the number of random genomes that were perturbed in this approach in each profile was the same as the number of genomes encoding the focal gene in the corresponding ‘focal gene’ profile. This set of simulated datasets will be referred to below as the ‘random taxa’ profiles. After producing these simulated profiles, we applied POMS, as well as the other differential abundance approaches to identify CEFs and differentially abundant functions, respectively.

We evaluated the obtained results by comparing the rank of the focal gene (based on p-value, from the smallest to the highest p-value) amongst all the significant functions identified by each approach. Under our focal gene-based simulations, MAGs encoding the focal genes across the profiles were the only direct targets of selection, and accordingly the focal gene was expected to be highly ranked (i.e., closer to 1, indicating a lower p-value compared to other genes) in the set of all significant genes.

Indeed, a drastic difference can be seen through ranking the resulting KOs based on these approaches (**Figure 2a**). Specifically, the focal genes in the POMS output are ranked significantly higher (median=1.5; mean=44.5; SD=104.8; P < 10^−15^) compared with the DESeq2 output (median=65.7; mean=291.7; SD=606.8), which was the next-highest-ranked output. This result highlights that genes conferring a selective advantage can in principle be better identified by POMS than standard differential abundance approaches. Overall, this finding was robust to the exact parameters used to simulate selection, and whether the simulations were based on MAGs or reference genomes, for these analyses (**see Supp. Results**; **Supp. Figures 2-6**).

**Figure 2:**
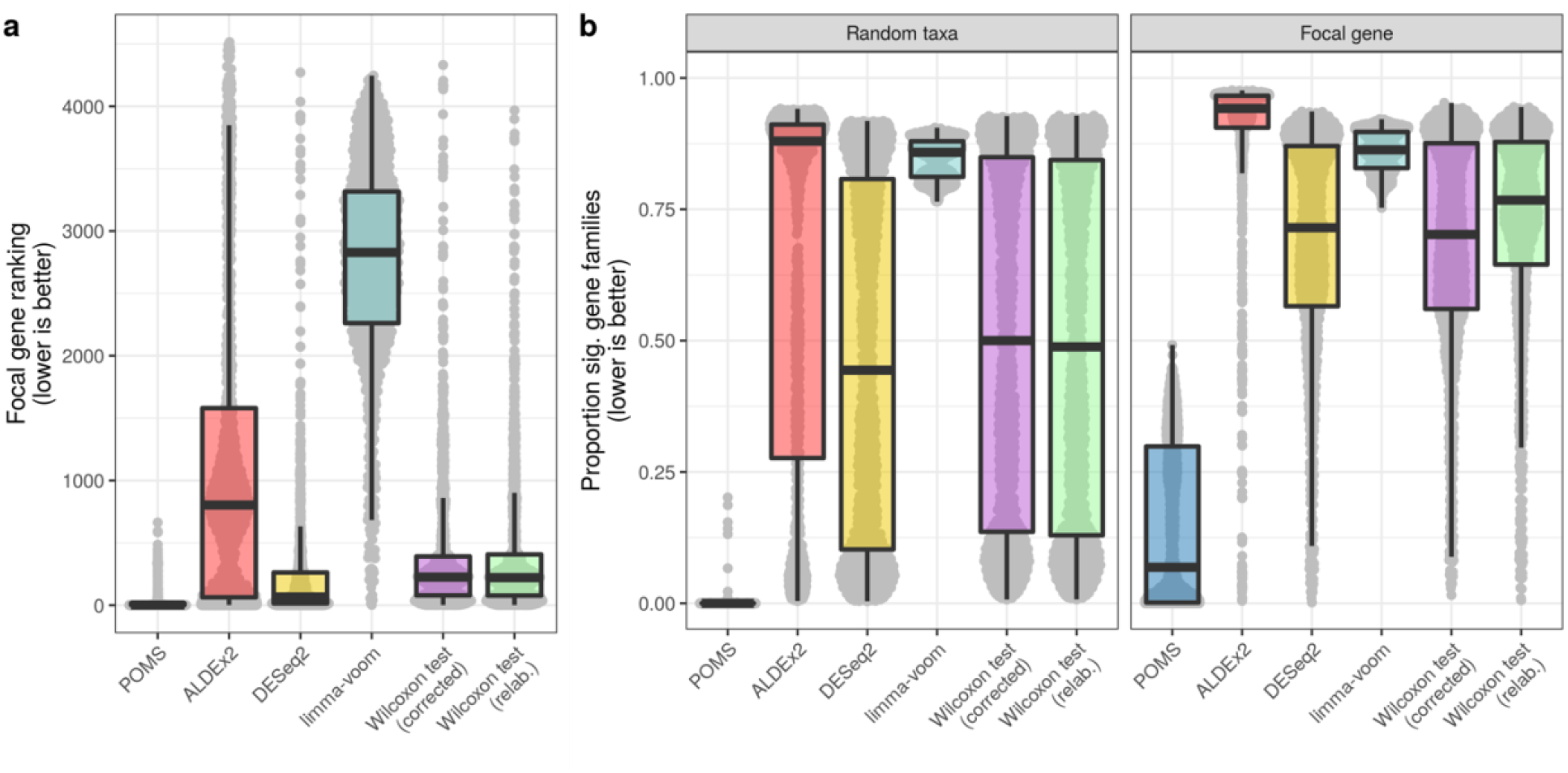
POMS performs best on simulated data based on focal gene rankings and the proportion of significant gene families. (a) Ranking of focal gene (i.e., the gene expected to be significant) in the output of all significant genes, based on p-values. (b) Proportion of gene families identified as significantly different between the simulated sample groups by each approach. The ‘Random taxa’ panel corresponds to the case where the abundance of randomly selected metagenome-assembled genomes was perturbed in one sample group. Under these conditions no systematic differences in functional abundances would be expected on average. The ‘Focal gene’ panel represents the simulations where the metagenome-assembled genomes that encode a specific gene family increased in abundance in one sample group. Each grey point corresponds to one simulation replicate profile. Note that “lower is better” is indicated on panel b to emphasize that identifying almost all functions as significant is not useful and that few genes are expected to be significant in the random taxa tests.

We next investigated what factors were underlying the variation in focal gene ranking across the simulation replicate profiles. We suspected that the number of MAGs that encoded each focal gene would markedly impact the detection ability. Indeed, we found that in the POMS results that the focal genes were ranked highly overall, except for those encoded by very few MAGs (**Supp. Figure 7**). Moreover, focal genes identified by POMS with relative rankings within the top ten ranked KOs were encoded by a median of 229.5 MAGs, whereas those not included in the top ten ranked KOs were encoded by a median of only 24 MAGs. This trend was interestingly reversed in the Wilcoxon test results, which we focused our comparison on as it is a representative, fast-running, and intuitive standard differential abundance approach. Specifically, focal genes ranked within the top ten KOs were encoded by a median of 10 MAGs only, whereas those not included in the top ten KOs were encoded by a median of 72 MAGs. In addition, we found that the focal gene was amongst the significant KOs in 67.4% and 99.0% of the POMS and Wilcoxon test outputs, respectively. Most occurrences where the focal gene was not significant in the POMS output corresponded to cases where the gene was encoded by either extremely few or almost all MAGs (**Supp. Figure 8**). These results reflect the fundamental distinction between POMS and standard differential abundance tests: POMS gives more weight to genes consistently enriched in independent taxonomic lineages, which can be most clearly detected for widely encoded (but not ubiquitous) genes. In contrast, bag-of-genes approaches, like the Wilcoxon test, do not consider taxa-function links and thus call genes as significantly different even when these genes are linked to only a small number of taxa.

Having shown that the focal gene is successfully detected and highly ranked by POMS, we next turned to compare the overall number of significantly different KOs identified by POMS vs standard differential abundance tests. This analysis demonstrated that the proportion of significantly different KOs in the random taxa simulated profiles was over 500-fold lower based on POMS (mean=0.00095; SD = 0.012) compared with the Wilcoxon test (USCG-corrected) results (mean: 0.496; SD=0.333; **Figure 2b**). However, importantly, the random taxa and focal gene simulated profiles yielded substantially different POMS results, whereas the difference between the two profiles was less pronounced using the standard differential abundance tests. Specifically, there was a 685% increase in the mean proportion of significant KOs detected by POMS in the focal gene-based dataset compared to the random taxa dataset, while the corresponding increase based on the Wilcoxon test (USCG-corrected) was only 148%. This suggests that POMS better controls the false positive rate in cases where no consistent functional differences are expected, but can still identify substantial numbers of significant hits when there is a true signal. In contrast, the standard differential abundance tests frequently identified more than 50% of functions as significant even in cases where sample group differences were based on random taxonomic perturbations.

We next applied POMS to several real metagenomics datasets. Unlike in our simulated datasets above, in these datasets we have no clear expectations regarding which functions would be identified as CEFs, and our evaluation is accordingly vulnerable to subjective biases in interpretation. Yet, as we argue below, many of the significant CEFs identified by POMS are reasonable given the presence of similar findings in the literature.

We first focused on MAGs assembled as part of the Tara Oceans project^15^. Since strong selection pressures may act upon microbes across environmental gradients in the ocean^16,17^, these metagenomics samples represent an appealing test case for POMS. Our analysis focused specifically on 93 samples with 642 MAGs annotated with KOs. Environmental factors associated with the water samples were also available, including temperature, salinity, oxygen levels, nitrate concentrations, phosphate concentrations, and nitrogen dioxide concentrations. We applied POMS to this dataset to detect microbial functions associated with each of these factors independently. We considered KEGG pathways and modules in addition to KOs for this analysis. These higher-level functions were computed for each MAG independently based on the KO annotations (see Methods). Because the environmental factor data is continuous rather than partitioned into discrete groups, we identified BSNs based on Spearman correlations between the balances at each node and the factor levels. The direction of each BSN was taken by the sign of the correlation coefficient. We also binned CEFs in this analysis as either stringent, intermediate, or lenient, based on BH cut-offs of 0.05, 0.15, and 0.25, respectively.

Our analysis across all environmental factors identified in total four CEFs below the intermediate cut-off and four additional CEFs below the lenient cut-off (**Table 1**). These CEFs were associated with either phosphate or mean salinity levels (for which there were 13 and 11 BSNs, respectively), and indeed some of the identified functions likely reflect selective actions of these two factors. For instance, a pathway linked to biofilm formation, ko05111, was found to be associated with higher phosphate levels, in agreement with previous findings of the non-trivial relationship between phosphorus levels and biofilm formation^18,19^. Carotenoid biosynthesis (ko00906) was also positively associated with phosphate levels, potentially reflecting a previously characterized and profiled phosphate-dependent pathway for carotenoid biosynthesis in marine bacteria^20^. Reasonable connections can also be made with the CEFs associated with mean salinity levels, such as the positive association with limonene and pinene degradation (ko00903). Limonene and pinene are monoterpenes that can be metabolized to modify cell membrane fluidity. This could reflect a potential adaptation that could facilitate organisms’ survival in the face of high salinity levels^21^.

**Table 1:** Tara Oceans POMS results.

| Env. factor | Data type | Func. id. | Total FSNs | FSNs ∩ BSNs (high)<sup>a</sup> | FSNs ∩ BSNs (low)<sup>b</sup> | FSNs not ∩ BSNs<sup>c</sup> | Cut-off | Corr. P-value | Function description |
| --- | --- | --- | --- | --- | --- | --- | --- | --- | --- |
| Phosphate | KO | K07137 | 11 | 9 | 0 | 2 | Inter. | 0.082 | Uncharacterized protein (COG2509: FAD-dependent dehydrogenase) |
| Phosphate | KO | K15523 | 9 | 8 | 0 | 1 | Inter. | 0.082 | FN3KRP; protein-ribulosamine 3-kinase [EC:2.7.1.172] |
| Mean salinity | Pathway | ko00903 | 15 | 8 | 0 | 7 | Inter. | 0.099 | Limonene and pinene degradation |
| Phosphate | Pathway | ko05111 | 13 | 8 | 0 | 5 | Inter. | 0.104 | Biofilm formation - *Vibrio cholerae* |
| Mean salinity | Module | M00535 | 11 | 7 | 0 | 4 | Len. | 0.196 | Isoleucine biosynthesis, pyruvate => 2-oxobutanoate |
| Phosphate | Pathway | ko00906 | 9 | 6 | 0 | 3 | Len. | 0.203 | Carotenoid biosynthesis |
| Phosphate | Module | M00044 | 12 | 8 | 0 | 4 | Len. | 0.210 | Tyrosine degradation, tyrosine => homogentisate |
| Mean salinity | Module | M00529 | 6 | 0 | 5 | 1 | Len. | 0.248 | Denitrification, nitrate => nitrogen |

<sup>a</sup>FSNs intersecting BSNs associated with higher levels of environmental factor
<sup>b</sup>FSNs intersecting BSNs associated with lower levels of environmental factor
<sup>c</sup>FSNs not intersecting BSNs

We next investigated how the results would differ if a standard approach for testing for associations between microbial functions and the environmental factors were applied instead. To this end, we computed Spearman correlations between every microbial function and every environmental factor. As expected, there were substantially more functions associated with each factor based on this approach. For instance, there were 3,955 and 4,916 significant KOs (BH < 0.25) associated with mean salinity and phosphate levels, respectively. All observed CEFs identified by POMS were also significant in the Spearman correlation output, but these functions were not consistently amongst the most highly ranked in this output (**Supp. Figure 9a**).

We next investigated how POMS performs on case-control metagenomics datasets. We focused on a dataset of MAGs compiled from human-associated microbiomes that were published as part of a large-scale meta-analysis^22^. We used subsets of this dataset corresponding to three disease datasets: two obesity datasets and one colorectal cancer dataset.

The first obesity dataset we analyzed included 477 obese and 257 control individuals that contained a total of 1,401 MAGs in their stool microbiomes. We applied POMS to this dataset and identified 34 BSNs that differed between obese and control individuals. The second obesity dataset analyzed included 251 obese and 159 control individuals that harboured a total of 1,161 MAGs in their stool microbiomes, with 31 resulting BSNs. Applying POMS, we identified 27, 3, and 23 consistently enriched KOs, pathways, and modules, respectively, in at least one dataset (**Supp. Table 1**). Of these hits, one pathway and four modules were called as CEFs across both datasets, but there were no overlapping KOs.

Many of these significant hits are reasonable given our knowledge of the link between the human microbiome and obesity. For example, the strongest CEF detected by POMS was for the module cytochrome bd ubiquinol oxidase (M00153), which was associated with case samples in the second obesity dataset. This function could reflect a broad shift in energy production through oxidative phosphorylation, or potential adaptation to oxidative stress, in obese individuals^23^. Biotin biosynthesis (M00123) was found by POMS to be associated with obesity in both datasets. Although biotin levels of obese individuals have not been well-studied, biotin deficiency has been associated with higher blood glucose levels and insulin resistance^24^. Beta-lactam resistance (ko01501) was also significantly associated with case samples in each dataset, which is particularly interesting as exposure to antibiotics has long been known to be associated with obesity^25^. One of the sole CEFs associated with control patients (and in the first obesity dataset only) was module M00545, which represents trans-cinnamate degradation to acetyl-CoA. This is also an intriguing observation as trans-cinnamate has been shown to be an effective treatment for obesity across several experimental conditions^26,27^.

Last, we applied POMS to the microbial profiles of stool samples from 75 colorectal cancer patients and 53 controls, which contained 1,187 MAGs. In this case, there were only 14 BSNs and the sole CEF that was detected by POMS was glyoxylate and dicarboxylate metabolism (enriched in control patients; corrected P=0.201). Interestingly, this has previously been identified as the most significantly depleted function in colorectal cancer samples (relative to controls) based on metabolite profiles^28^.

Similar to the Tara Oceans dataset, the CEFs identified by POMS in these human disease datasets were almost all significant based on standard differential abundance approaches. However, once again these functions were not amongst the top hits identified by these approaches overall (**Supp. Figure 9b-d**).

## Discussion

Herein we have presented and validated the POMS framework: a novel approach for identifying consistently enriched functions in microbiome data. Using simulations, we have demonstrated that POMS can accurately identify widely encoded microbial functions that confer a strong selective advantage. While we present several different analyses to validate and justify our approach, perhaps the most convincing evidence is that focal genes (i.e., genes that were simulated as conferring a selective advantage) were amongst the most significant functions identified by POMS, whereas this was often not the case for standard differential abundance approaches. POMS also produced sensible results on real shotgun metagenomics datasets that were assembled into MAGs, as many of the identified CEFs agree with known findings from the literature.

Although our validation analyses of POMS are promising, our approach has several caveats. First, POMS can only identify functions that are widely encoded by taxa and that are variably present across different lineages; a gene restricted to an individual lineage, even if that lineage has many members, may not be identified as a CEF. This is because POMS requires a function to be enriched repeatedly (and in consistent directions) at independent BSNs to identify a function as significant. Consequently, POMS may miss many functions that would otherwise be identified by more standard approaches. However, CEFs identified by POMS are easier to interpret: each CEF represents the hypothesis that the function provides a fitness advantage in certain contexts to those taxa which encode it. Thus, we believe that the general approach used by POMS is not a replacement for standard differential abundance tests, but rather a proof-of-concept that alternative approaches can well-complement existing methods.

It should also be appreciated that significant CEFs identified by POMS are not guaranteed to be enriched only in a single direction. The multinomial test executed by POMS tests for whether the distribution of FSNs into the three categories departs from the random expectation. Significant functions could simply be enriched at BSNs of both types, i.e., a mixture of BSNs relatively higher in both sample groups. Such cases could still be biologically interesting but would be interpreted differently than CEFs primarily enriched towards a single sample group. Accordingly, the counts of FSNs of each category should be considered when interpreting any CEFs identified by POMS.

There are also other limitations to the POMS approach. For instance, identifying CEFs requires that sufficient BSNs are present to identify significant enrichments. Even if an adaptive function is widely distributed phylogenetically, it will not be identifiable by POMS unless there are corresponding BSNs that could be driven by this function. In addition, POMS assumes that all nodes in the tree have independent balance distributions. This is partially invalid because different taxonomic groups are more likely to vary across individuals and taxa co-occurrence can occur even at long evolutionary distances^29^.

Despite these caveats, we have shown that compared with more common approaches, integrating functional information into a balance tree framework can better identify functions that could provide a selective advantage. POMS is just one example of how this general analysis scheme can be implemented, and future work could incorporate more sophisticated approaches for analyzing balances across phylogenetic trees^8,30^. Nonetheless, the current POMS framework represents a more robust methodology compared with existing statistical tests applied in the microbiome field. Given the wide range of results across microbiome differential abundance approaches^31^, clearly more reliable approaches are needed, even if they come at the cost of lower sensitivity.

## Methods

### Isometric Log-Ratio

Phylogenetic balances in POMS are calculated based on the isometric log-ratio of taxa in the two node subtrees (i.e., on one side of the node compared to the other)^8,9^. Specifically, the balance for a sample at node *i* is calculated based on the equation

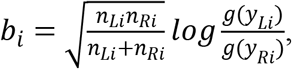

where *n*_*Li*_ and *n*_*Ri*_ correspond to the numbers of taxa on the left- and right-hand sides of the node. Similarly, *g*(*y*_*Li*_) and *g*(*y*_*Ri*_) correspond to the geometric means of the relative abundances of taxa on the left- and right-hand sides of the node. Note that the choice of which lineage is considered the left-versus right-hand side of a given node is arbitrary. This approach converts microbiome relative abundance data into ratios of geometric means. The numbers of taxa on each side of a node are included in this calculation to scale the balance to give it unit length (i.e., to make the balances comparable despite varying numbers of taxa at each node).

The geometric mean of the relative abundance of a set of taxa on the left-hand side is calculated based on the equation

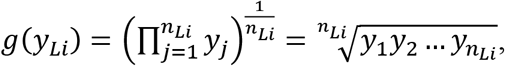

and analogously for the right-hand side.

### Multinomial test

A multinomial exact test is used in POMS to identify CEFs. This test considers the counts of three classes of FSNs per function: (1) FSNs that do not intersect with BSNs, (2) FSNs that intersect with BSNs where the functional enrichment is in taxa that are relatively more abundant in sample group one, and (3) FSNs that intersect with BSNs where the functional enrichment is in taxa that are relatively more abundant in sample group two. The null expectation for the proportion of FSNs in each category is based on a mass-action interaction between FSNs and BSNs. In other words, the null expectation corresponds to the case where FSNs and BSNs are assigned randomly. The expectation also assumes that FSNs that intersect with BSNs are split equally between BSNs that are higher in sample groups one and two, respectively. This test is implemented with the xmulti function of the XNomial R package^32^ (tested with version 1.0.4).

### Metagenome-assembled genome-based simulations

The MAG-based simulations were based on 704 control samples from a large human meta-analysis dataset^22^. These samples all met the metadata criteria of being labelled as healthy, adult, and not on antibiotics. These samples were also from studies that included at least 40 samples in total. These simulations proceeded as described in the Results section. First, the samples were randomly split into two groups for each of the 1,000 replicate datasets. Then, for each profile a focal gene family was randomly chosen, and within one group the abundance of all MAGs encoding this gene family was incremented by a pseudocount of 1 and these abundances were multiplied by a factor of 1.5. These simulated datasets were then input to POMS and the results are referred to as the “focal gene” profiles. We also performed parallel simulations where the relative abundance of random taxa was randomly inflated by identical amounts in one group only. Importantly, the same number of taxa were perturbed as were affected in each matching focal gene simulation profile. The resulting output based on these simulated profiles are referred to as the “random taxa” profiles.

The alternative differential abundance approaches compared in this study were: Wilcoxon tests following normalization by the median USCG abundance, using the identical approach used by the tool MUSiCC^33^, Wilcoxon tests based on relative abundances, ALDEx2 v1.16.0, DESeq2 v1.24.0, and limma-voom v3.40.6. Significant gene families were identified based on a BH cut-off < 0.05 for all tested approaches, including POMS.

### Tara Oceans dataset validation

The assembled Tara Oceans metagenomics dataset was taken from a pre-existing project^15^. The original 957 MAGs were downloaded from FigShare (https://figshare.com/articles/dataset/TARA-NON-REDUNDANT-MAGs/4902923). Environmental and chemical profiles of the ocean samples along with the relative abundance and annotations of MAGs were taken from the published supplementary tables for this project^15^.

GToTree^34^ v1.4.16 was run to build a phylogenetic tree based on shared single-copy genes and to exclude MAGs with completeness below 60% and redundancy above 10%, which resulted in retaining 642 MAGs. MAGs were called as present within samples if the breadth of coverage was > 1%. This is a lenient setting, which was chosen based on the empirical distributions of MAG breadth of coverage across the samples.

Higher-level functions (i.e., KEGG modules and pathways) were reconstructed based on KEGG mappings from KOs to these categories downloaded from the KEGG website^10^ on April 12^th^, 2021. Reconstruction was performed using the PICRUSt2 script pathway_pipeline.py^35^, which leverages MinPath^36^ and the algorithm implemented in HUMAnN2^37^ to reconstruct higher-level function abundances. These reconstructions were performed for each MAG independently.

Because the environmental data, such as the salinity and nutrient concentrations, were continuous, significant BSNs were identified based on Spearman correlations between sample balances and these environmental factors. These results were then discretized to indicate whether the factors were positively or negatively associated with the groups at each respective BSN.

### Case-control shotgun metagenomics dataset validations

We focused our validation analyses on three datasets that were part of the large meta-analysis of human shotgun metagenomics datasets^22^. These datasets are defined based on dataset accession identifiers in the European Nucleotide Archive. The largest dataset focused on obese and control individuals (which we refer to as the first obesity dataset) and corresponds to data accession identifier ERP002061. The second obesity dataset corresponds to accession identifier ERP003612 and the colorectal cancer dataset is under accession identifier ERP012177. We used the previously generated MAGs, sample MAG abundance profiles, and MAG phylogenetic tree as input to POMS after performing pre-processing. Importantly, we excluded MAGs from each sample with mapped read coverage lower than 25%. KEGG pathways and modules were reconstructed as for the Tara Oceans dataset.

### POMS Dependencies

POMS is written in R^38^ and is dependent on the following R packages (versions used in this manuscript are indicated, but these exact versions are not required): ape^39^ v5.3, parallel v3.6.0, phangorn^40^ v2.5.5, and stringr^41^ v1.4.0. Testing and development of this approach was carried out using R v3.6.0 and RStudio v1.2.5033 on a server running Ubuntu v16.04.5.

Several additional R packages are required to follow the current analysis workflow after running POMS (again the versions indicated were used for this paper, but are not required versions): ggtree^42,43^ v1.16.1, ggplot2^44^ v3.3.0, plyr^45^ v1.8.4, and reshape2^46^ v1.4.3. All multi-panel plots displayed were created with the cowplot^47^ (v1.0.0) R package.

## Code Availability

The code for all analyses presented in this manuscript is available at: https://github.com/gavinmdouglas/POMS_manuscript/. The source code for POMS is available at: https://github.com/gavinmdouglas/POMS.

## Acknowledgements

We would like to thank Cecilia Noecker for helpful feedback on POMS. GMD was funded by a National Sciences and Engineering Research Council (NSERC) Canada Graduate Scholarship (Doctoral). MGIL is supported by an NSERC Discovery Grant. EB is a Faculty Fellow of the Edmond J. Safra Center for Bioinformatics at Tel Aviv University.

