## Supplementary Materials for "Integrating phylogenetic and functional data in microbiome studies"

### <u>Supplementary Methods and Results</u>

#### Exploring the parameter space of the MAG-based simulations

The selection simulation framework that we describe in the main text represents an extremely strong selection acting upon taxa encoding the focal gene in each profile. Specifically, this simulated selective advantage included (in the sample group experiencing selection) adding a pseudocount of one to the abundance of all taxa that encoded the focal gene and then multiplying the resulting abundance by 1.5.

To confirm that POMS is not only performing better under this extreme condition we also created and tested simulated profiles over a range of weaker selection settings. These altered settings included combinations of not adding a pseudocount and multiplying the abundances by lower factors (1.05, 1.1, and 1.3). We additionally tested the impact of different numbers of metagenome-assembled genomes (MAGs) in these simulations, to investigate how dataset size affects the POMS framework. This analysis clearly demonstrated that across most settings the focal genes were substantially more highly ranked by the POMS framework compared with the Wilcoxon test approach (**Supp. Figure 2**).

The exception was for simulation settings where no pseudocount was added, where the focal gene was largely non-significant in the POMS output due to insufficient statistical power. This drop in statistical power is reflected by lower numbers of balance-significant nodes (BSNs) on average for these settings (**Supp. Figure 3**). In contrast, the focal genes were highly ranked on average based on the Wilcoxon test for these simulation settings, which is consistent with our finding that the Wilcoxon test identifies functions as more highly ranked when they are encoded by a small number of taxa (**Supp. Figure 7**).

Our analysis also demonstrated a marked drop in the number of BSNs in simulations in simulations with fewer than 250 MAGs (**Supp. Figure 3**). POMS was unable to call any focal genes as significant in these simulations with lower sample sizes due to the small number of BSNs, while many were called as significant by the Wilcoxon test.

Finally, these analyses also demonstrated that the overall trend of the Wilcoxon test identifying a higher proportion of significant hits relative to POMS was robust to the simulation settings (**Supp. Figure 4**).

**Reference-genome-based simulations**

Our observations based on the MAG-based simulations are valuable, but one caveat is that the quality of published MAGs is often questionable[1]. To ensure that misassembled MAGs were not driving our results, we repeated our simulation approach on 500 reference genomes.

These genomes were taken from the Integrated Microbial Genomes database[2] that were previously parsed for use with PICRUSt2[3]. Per-genome KEGG ortholog annotations were taken from the default PICRUSt2 database. We created a de novo phylogenetic tree based on a set of universal single-copy genes with GToTree[4] v1.4.16. This approach parses out universal single-copy genes from genome sequences and wraps several tools to build a phylogenetic tree. The tool was run with the bacterial hidden Markov model setting and with FastTree[5] v2.1.10. GToTree also returns estimates of the percent completeness and redundancy for each genome. We excluded all genomes with completeness below 95% and/or redundancy above 5%. We then randomly sampled 500 of the remaining high-quality genomes for the subsequent analyses.

We next simulated random abundances of these genomes across 1,000 samples based on the zero-inflated beta distribution implemented in the rBEZI function of the gamlss.dist v5.1.7 R package[6]. Simulations under this model can be modified with three key metrics: mu (the mean), nu (the probability of zero abundance), and sigma (the standard deviation). For these simulations we maintained values of mu and sigma of 0.1 and 1, respectively, throughout. We generated four simulated datasets based on nu values set to 0.5, 0.65, 0.8, and 0.95. When generating these simulated datasets we required that each sample contain a minimum of five genomes (i.e., simulated profiles were re-run if fewer than five genomes had non-zero abundance). These altered nu values had a large impact on the sparsity and inter-sample overlap of each dataset (**Supp. Figure 5a**).

We than re-ran the key steps of our simulation analysis on these simulated abundance tables and genomes. Specifically, we repeated the focal gene-based simulation, including applying POMS and Wilcoxon tests, over 500 replicate profiles for each of the four datasets. The focal gene ranks varied substantially depending on the abundance table simulation approach (**Supp. Figure 5b**). In addition, the focal gene ranks were significantly higher in the POMS output compared with the Wilcoxon test output ($P < 10^{-15}$), with one exception (nu=0.5; W=18,416; P=0.157). This analysis confirmed that the key findings reported in the main text, and the ability of POMS to pinpoint selected functions, carry over to these new datasets and are not an artifact of using MAGs in our prior simulations.

We also investigated the relationship between focal gene ranks and the number of genomes encoding the focal gene. Like in the MAG-based simulation results, those significant focal genes in the POMS output that were not amongst the most significant genes (i.e., focal genes at higher ranks) were encoded by fewer genomes (**Supp. Figure 6**). In contrast, the focal gene ranks based on Wilcoxon test p-values displayed a positive linear relationship with the number of encoding genomes. The strength of this relationship increased with the sparsity of the data, and indeed the sparsest simulated dataset (nu=0.95) showed high correlation (Pearson $R=0.876$; $P < 10^{-15}$). This result was similar to the positive correlation between focal gene rank and number of encoding genomes that we also observed in the corresponding analysis on the MAG-based simulations (**Supp. Figure 7a**). Overall, these reference genome-based simulation results are consistent with the key observations from the MAG-based simulations. In addition, the reference genome simulations based on tables of varying sparsity highlight how POMS performs better with highly sparse data, which is characteristic of microbiome datasets.

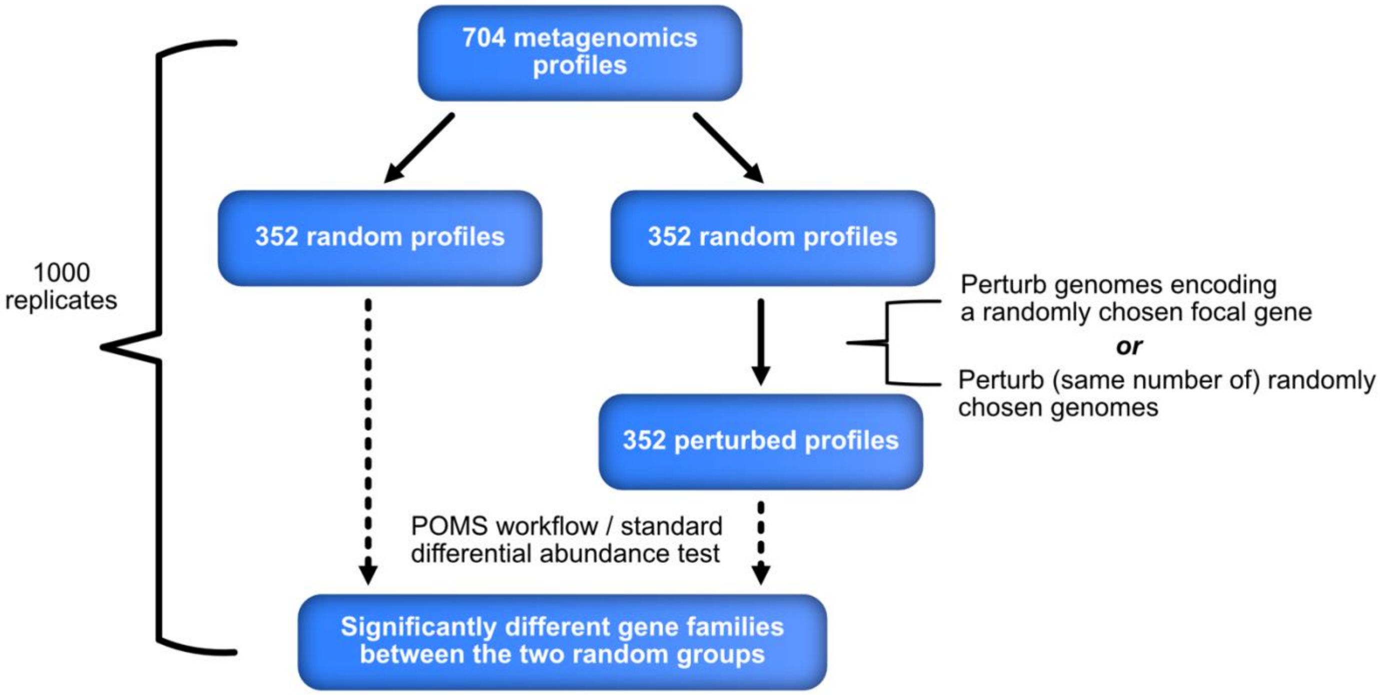


**Supplementary Figure 1: Workflow diagram for metagenome-assembled genome-based simulations**.

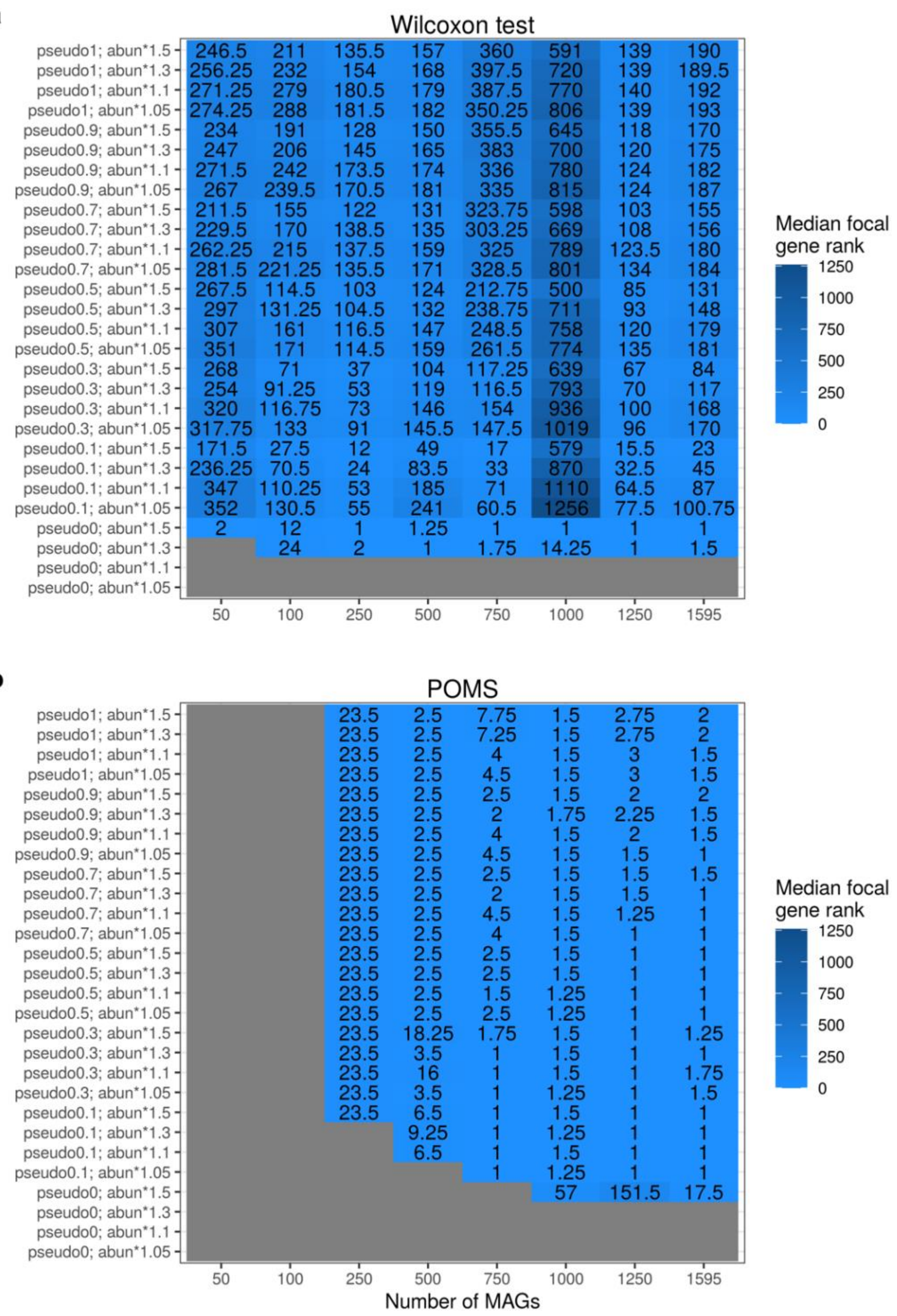


**Supplementary Figure 2: Median ranking of focal gene in output of (a) Wilcoxon test and (b) POMS across replicates per simulation setting.** The “pseudo” setting is the proportion of metagenome-assembled genomes that encoded the focal gene that were randomly selected per sample to be given a pseudocount of 1 to their abundance. The “abun” setting represents the scaling factor of the abundance of each genome encoding the focal gene after this pseudocount step. Grey boxes indicate cases where the focal gene was not significant in any replicate. MAGs: metagenome-assembled genomes.

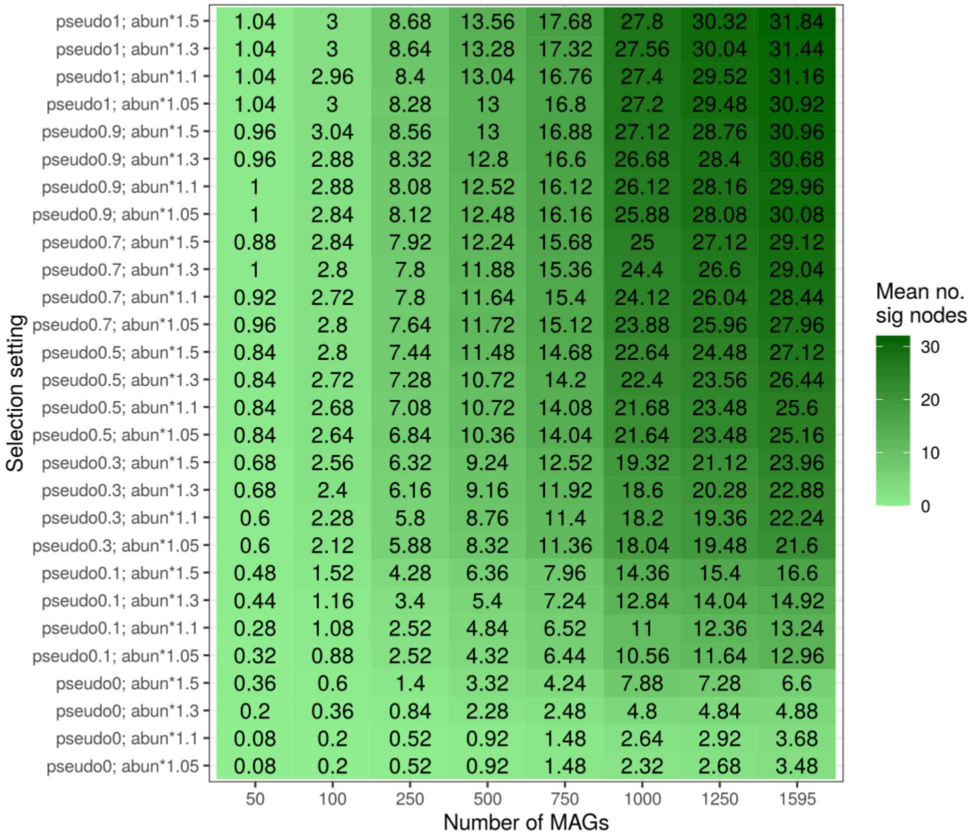


**Supplementary Figure 3: Mean number of significant nodes based on sample balances (i.e., balance-significant nodes) across replicates for each simulation setting.** The "pseudo" setting is the proportion of metagenome-assembled genomes that encoded the focal gene that were randomly selected per sample to be given a pseudocount of 1 to their abundance. The "abun" setting represents the scaling factor of the abundance of each genome encoding the focal gene after this pseudocount step. MAGs: metagenome-assembled genomes.

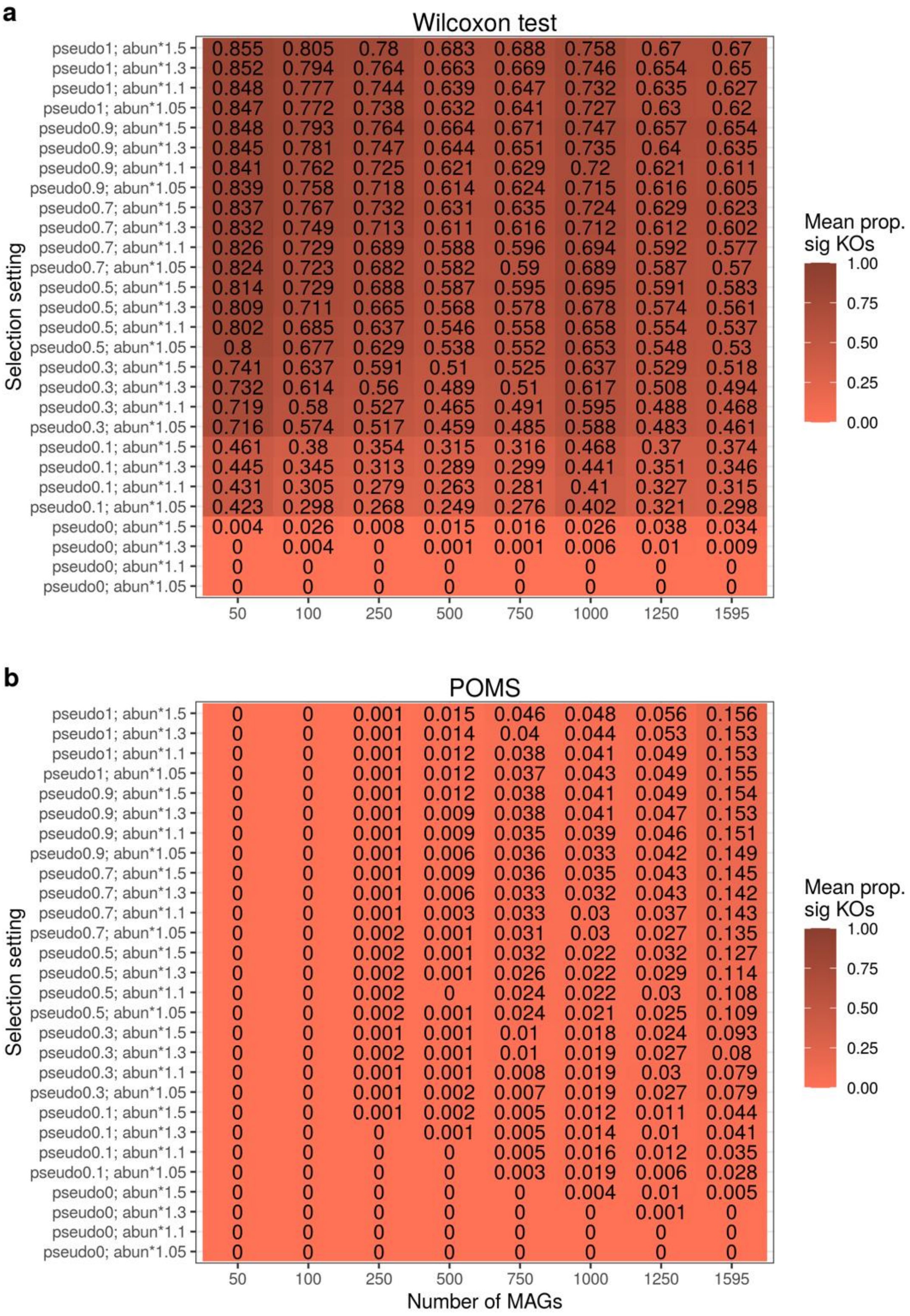


**Supplementary Figure 4: Mean proportion of significant functions in output of (a) Wilcoxon test and (b) POMS across focal gene-based replicates per simulation setting.** The "pseudo" setting is the proportion of metagenome-assembled genomes that encoded the focal gene that were randomly selected per sample to be given a pseudocount of 1 to their abundance. The "abun" setting represents the scaling factor of the abundance of each genome encoding the focal gene after this pseudocount step. MAGs: metagenome-assembled genomes; KOs: KEGG orthologs (i.e., the tested functions).

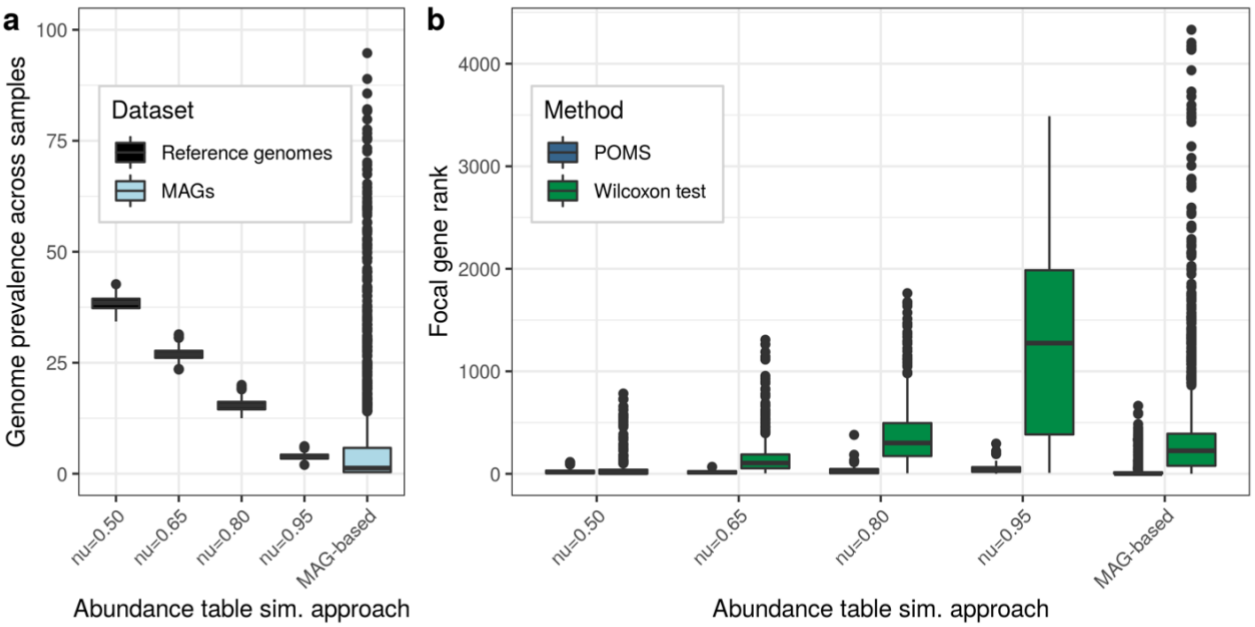


**Supplementary Figure 5: Genome prevalence and ranking percentiles of focal genes varies across all simulation datasets.** (a) The prevalence (%) of each genome (or metagenome-assembled genome [MAG]) across all samples in a dataset. (b) The ranking percentiles of the focal gene within the list of significant genes for each simulated dataset setting. The reference genome-based simulated datasets were altered based on four parameter settings, which greatly affect genome prevalence. The “MAG-based” group corresponds to the simulation results shown in the main text. This category is displayed to enable clear comparisons with the MAG-based simulation results reported in the previous section.

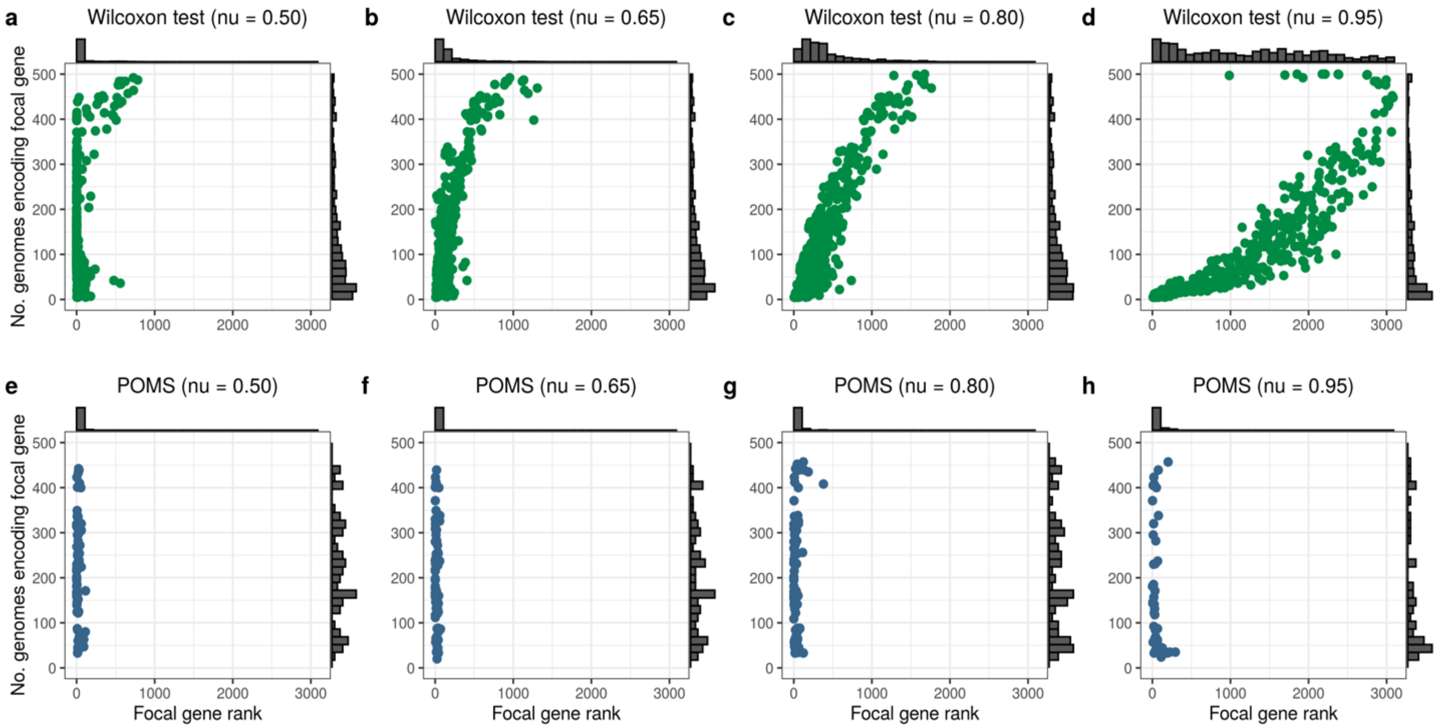


**Supplementary Figure 6: Ranking percentiles of focal genes against the number of genomes in which they are encoded, based on reference genome simulations.** Results based on the Wilcoxon test (universal single-copy gene abundance corrected) are shown in green and those based on POMS in blue. Simulation setting (determined by the value of nu, the probability that a feature has an abundance of 0) is indicated above each panel. The grey histograms represent the marginal distributions.

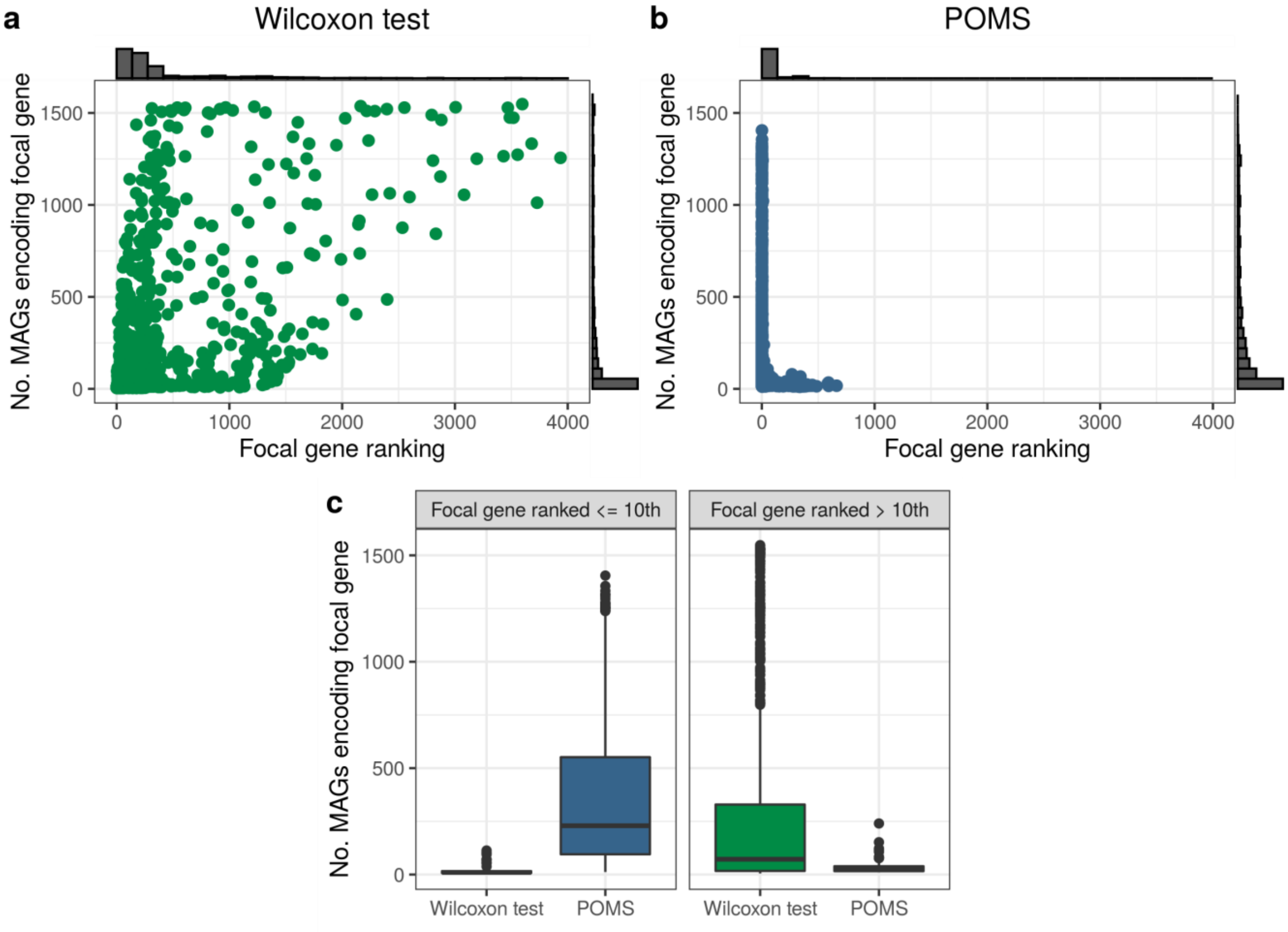


**Supplementary Figure 7: Focal gene rankings are strongly associated with the number of metagenome-assembled genomes (MAGs) in which the focal gene is encoded.** Each point is a simulation replicate analyzed with (a) the Wilcoxon test approach or (b) POMS. The grey histograms represent the marginal distributions. (c) Number of MAGs that encode the focal gene, plotted separately for those ranked within and outside the top ten significant hits.

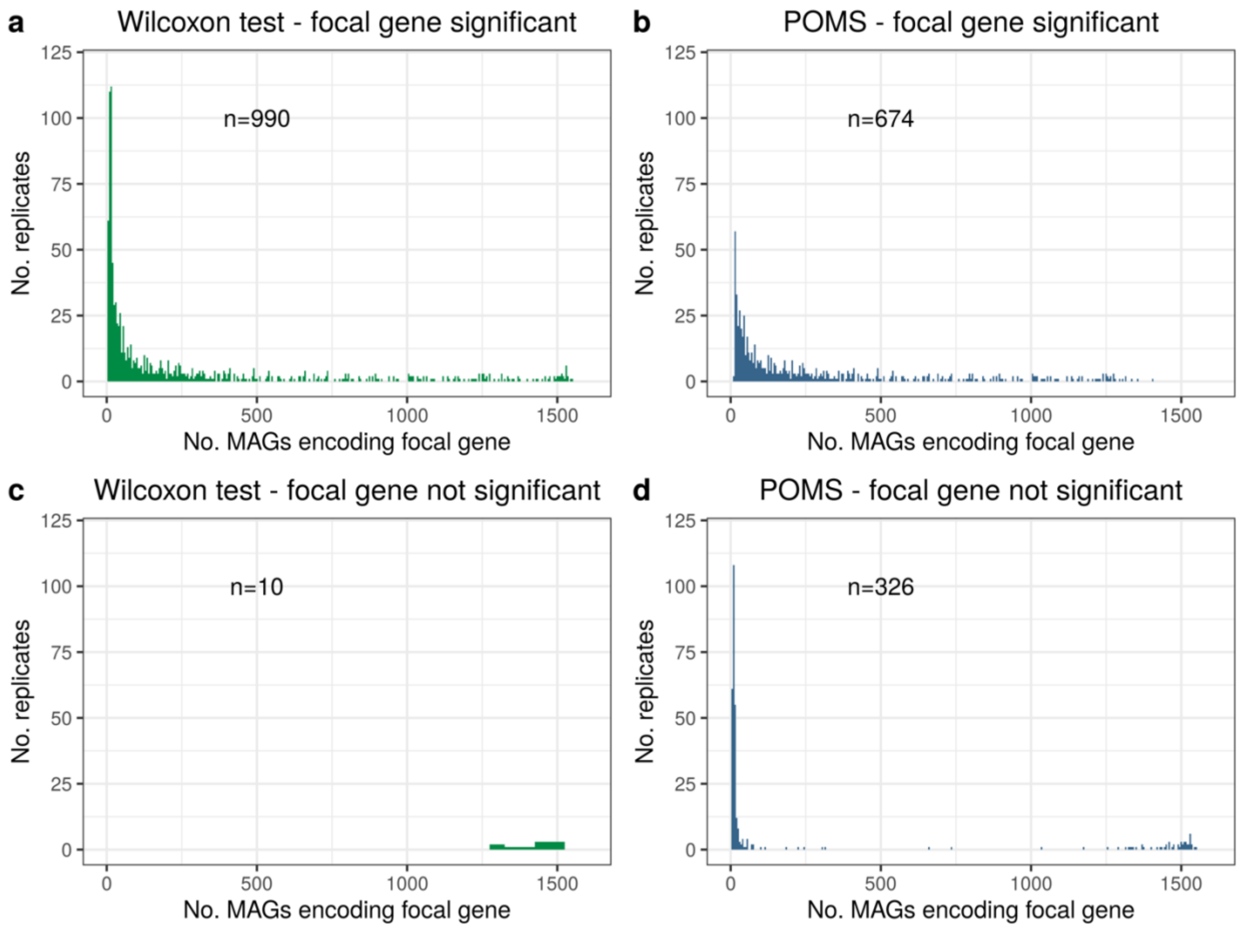


**Supplementary Figure 8: Number of metagenome-assembled genomes (MAGs) that encode the focal gene in cases where it was called either significant or not.** Number of replicates shown in each case is indicated within each panel.

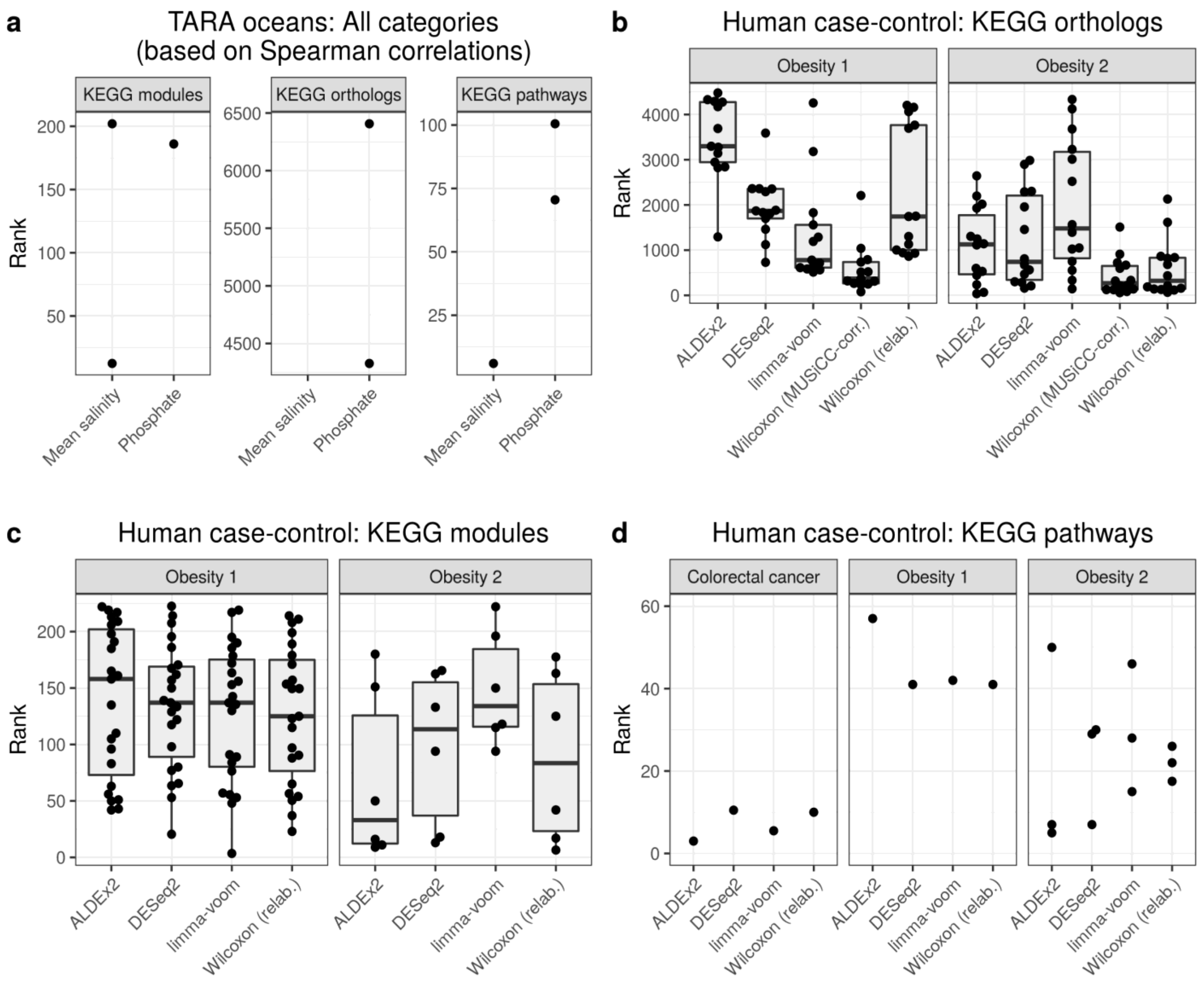


**Supplementary Figure 9: Consistently enriched functions (CEFs) in POMS output are not generally amongst the top significant hits based on standard statistical approaches.** Each dot corresponds to the rank of a CEF (i.e., a function identified as significant based on POMS) in the list of significant functions identified by an alternative tool.

**Supplementary Table 1: Significant KEGG functions in obesity datasets based on POMS (only showing hits with corrected P-value < 0.2)**

| Dataset | Func. Type | Function | Corr. P-value | Description |
|---|---|---|---|---|
| Obesity 2 | Module | M00153 | 0.0099 | Cytochrome bd ubiquinol oxidase |
| Obesity 2 | KO | K01993 | 0.0493 | ABC-2.TX; HlyD family secretion protein |
| Obesity 2 | Module | M00123 | 0.082 | Biotin biosynthesis, pimeloyl-ACP/CoA => biotin |
| Obesity 2 | Module | M00082 | 0.082 | Fatty acid biosynthesis, initiation |
| Obesity 2 | Module | M00116 | 0.082 | Menaquinone biosynthesis, chorismate (+ polyprenyl-PP) => menaquinol |
| Obesity 2 | KO | K00425 | 0.0903 | cydA; cytochrome bd ubiquinol oxidase subunit I [EC:7.1.1.7] |
| Obesity 2 | KO | K02523 | 0.0903 | ispB; octaprenyl-diphosphate synthase [EC:2.5.1.90] |
| Obesity 1 | KO | K09861 | 0.1208 | K09861; uncharacterized protein |
| Obesity 1 | KO | K00941 | 0.1208 | thiD; hydroxymethylpyrimidine/phosphomethylpyrimidine kinase [EC:2.7.1.49 2.7.4.7] |
| Obesity 1 | Module | M00123 | 0.1425 | Biotin biosynthesis, pimeloyl-ACP/CoA => biotin |
| Obesity 1 | Module | M00048 | 0.1425 | Inosine monophosphate biosynthesis, PRPP + glutamine => IMP |
| Obesity 1 | Module | M00880 | 0.1425 | Molybdenum cofactor biosynthesis, GTP => molybdenum cofactor |
| Obesity 1 | Module | M00015 | 0.1425 | Proline biosynthesis, glutamate => proline |
| Obesity 1 | KO | K01923 | 0.143 | purC; phosphoribosylaminoimidazole-succinocarboxamide synthase [EC:6.3.2.6] |
| Obesity 1 | KO | K01933 | 0.143 | purM; phosphoribosylformylglycinamidine cyclo-ligase [EC:6.3.3.1] |
| Obesity 1 | Pathway | ko01501 | 0.1514 | beta-Lactam resistance |
| Obesity 2 | Pathway | ko01501 | 0.1571 | beta-Lactam resistance |
| Obesity 2 | Pathway | ko00540 | 0.1571 | Lipopolysaccharide biosynthesis |
| Obesity 1 | Module | M00525 | 0.1575 | Lysine biosynthesis, acetyl-DAP pathway, aspartate => lysine |
| Obesity 1 | Module | M00526 | 0.1575 | Lysine biosynthesis, DAP dehydrogenase pathway, aspartate => lysine |
| Obesity 1 | Module | M00116 | 0.1589 | Menaquinone biosynthesis, chorismate (+ polyprenyl-PP) => menaquinol |
| Obesity 1 | KO | K02500 | 0.1592 | hisF; imidazole glycerol-phosphate synthase subunit HisF [EC:4.3.2.10] |
| Obesity 1 | KO | K01952 | 0.1592 | PFAS, purL; phosphoribosylformylglycinamidine synthase [EC:6.3.5.3] |
| Obesity 1 | KO | K01588 | 0.1592 | purE; 5-(carboxyamino)imidazole ribonucleotide mutase [EC:5.4.99.18] |
| Obesity 1 | Module | M00845 | 0.1656 | Arginine biosynthesis, glutamate => acetylcitrulline => arginine |
| Obesity 1 | Module | M00844 | 0.1656 | Arginine biosynthesis, ornithine => arginine |
| Obesity 1 | Module | M00793 | 0.1656 | dTDP-L-rhamnose biosynthesis |
| Obesity 1 | Module | M00082 | 0.1656 | Fatty acid biosynthesis, initiation |
| Obesity 1 | Module | M00119 | 0.1656 | Pantothenate biosynthesis, valine/L-aspartate => pantothenate |
| Obesity 1 | Module | M00899 | 0.1656 | Thiamine salvage pathway, HMP/HET => TMP |
| Obesity 1 | Module | M00545*** | 0.1656 | Trans-cinnamate degradation, trans-cinnamate => acetyl-CoA |
| Obesity 1 | Module | M00134 | 0.1701 | Polyamine biosynthesis, arginine => ornithine => putrescine |
| Obesity 1 | KO | K13038 | 0.1743 | coaBC, dfp; phosphopantothenoylcysteine decarboxylase / phosphopantothenate---cysteine ligase [EC:4.1.1.36 6.3.2.5] |

***This was the only function enriched towards control samples.
